# Kafka interfaces for composable streaming genomics pipelines

**DOI:** 10.1101/182030

**Authors:** Francesco Versaci, Luca Pireddu, Gianluigi Zanetti

## Abstract

Modern sequencing machines produce order of a terabyte of data per day, which need subsequently to go through a complex processing pipeline. The standard workflow begins with a few independent, shared-memory tools, which communicate by means of intermediate files. Given the constant increase of the amount of data produced, this approach is proving more and more unmanageable, due to its lack of robustness and scalability.

In this work we propose the adoption of stream computing to simplify the genomic pipeline, boost its performance and improve its fault-tolerance. We decompose the first steps of the genomic processing in two distinct and specialized modules (preprocessing and alignment) and we loosely compose them via communication through Kafka streams, in order to allow for easy composability and integration in the already existing Hadoop-based pipelines. The proposed solution is then experimentally validated on real data and shown to scale almost linearly.

## I. INTRODUCTION

Life sciences are quickly becoming one of the most prominent example of data driven science [1]–[3]. Next-generation sequencing (NGS) machines are among the main drivers of this phenomenon. They have brought huge improvements in DNA sequencing capacity together with drastic cost reductions, thus paving the way to a myriad of new applications that were previously technologically or economically unfeasible [4]. High-throughput sequencing can now be used for research into understanding human genetic diseases [5], in oncology [6], to study human phylogeny [7], and it is quickly becoming common in many other life sciences research sectors [8]. At the same time, NGS is reaching the level of personalized diagnostic applications [9], [10] and its continuous improvements in capacity are enabling very large scale population studies [11]. The output of a NGS machine is, basically, the digitalization of the results of a sequence of massively parallel biochemistry experiments and converting the raw data obtained to biologically relevant information requires to go through various intense processing steps. Thus, one of the main challenges that needs to be confronted is to develop scalable computing tools that can keep up with such a massive data generation throughput. The standard computational workflows start from the raw data produced by the NGS machines and are based on the use of shared-memory tools, which communicate by means of intermediate files. Traditionally [12], the workflow steps are run on a conventional High-Performance Computing (HPC) infrastructure – a set of computing nodes accessed through a batch queuing system and equipped with a parallel shared storage system. While this is, of course, a working solution, it requires a non–trivial amount of ad-hoc manual intervention to efficiently use the available computational resources and obtain the fast turn-around times that are needed for diagnostic applications [13]. The main issues here are how to divide the work of a single job among all computing nodes and how to make the system robust to transient or permanent hardware or software failures. Even though these issues can be surmounted [12], the dependency of the HPC infrastructure on a central shared storage system imposes an intrinsic limitation on the scalability of the platform. In fact, since some of the computational steps are I/O limited, access to storage can quickly become a bottleneck with the increase of the number of computational nodes.

As an alternative approach, there has been a recent interest [13]–[17] in supporting the NGS data processing pipelines using completely distributed platform such as Apache Hadoop. In this approach, the raw and intermediate data files are stored on a scalable file system, HDFS, and the computation is based on processing steps expressed, e.g., as Hadoop MapReduce, Spark or Flink programs. While this approach solves the I/O scalability issues and intrinsically provides resilience with respect to node failures, it is still based on the sequential application of the computational steps.

In this paper, we experiment the adoption of stream computing to help simplifying and boosting the performance of the genomic pipeline. Specifically, we decompose the first steps of the genomic processing in two distinct and specialized modules (preprocessing and alignment) and we loosely compose them via communication through Kafka streams, in order to allow for easy composability and integration in the already existing Hadoop-based pipelines. To the best of the authors’ knowledge, this is the first solution that can process the sequencer’s raw data using a fully scalable streaming approach and it is the first example of using a distributed publish-subscribe messaging system (Kafka) to decouple between processing steps of a bioinformatics pipeline.

The rest of the paper is organized as follows. Section II describes the Next Generation Sequencing process, the computation that is required to make sense of the data and the state-of-the-art in modern sequencing centers. Section III provides background regarding Flink and Kafka, motivating the decision to adopt them in our framework. Section IV present the tools that have been developed as part of this work. Section V contains the performance evaluation and a comparison to the state-of-the-art. Finally, Section VII discusses related work and Section VIII concludes the manuscript.

## II. THE NGS PROCESS

**Note:** This expository section is reproduced almost verbatim from [15], for the reader’s convenience.

Deoxyribonucleic acid (DNA) is a polymer composed of simpler units known as nucleotides, or *bases*. These come in four kinds: Adenine, Cytosine, Guanine and Thymine – respectively denoted by their initial letters A, C, G, and T. The DNA data produced by the NGS process are not directly interpretable from a biological point of view. In fact, the various high-throughput NGS technologies [18] all use a “shotgun” approach, where the genome to be sequenced is broken up into fragments of approximately the same size and the individual fragments are sequenced in parallel. The characteristics of the raw data produced by the sequencer changes depending on the specific technology being used.

The sequencers by Illumina Inc. (http://www.illumina.com) which are the target of this work – operate by successively attaching a fluorescent identifying molecule to each base of the DNA fragments being sequenced [19]: the genomic material is placed in a flowcell, that is organized in lanes which are, in turn, subdivided in tiles. At each cycle, the machine acquires a single base from all the reads by snapping a picture of the flowcell, where a chemiluminescent reaction is taking place. For each picture, and thus cycle, the software on the sequencer’s controlling workstation performs base calling from the image data – mapping each chemiluminescent dot to a specific base (A, C, G, T) based on its color. The process produces base call files (BCL), which contain the bases that were acquired from all the fragments – also known as *reads* but only in that specific sequencing cycle. Therefore, the individual DNA fragments are actually split over *C* files, where *C* is the number of cycles in the sequencing run and the length of the reads.

Next-generation sequencing activity quickly results in a lot of data. Consider that a modern high-throughput sequencer like the Illumina HiSeq 4000 can produce 1500 Gigabases, which equate to about 4 Terabytes of uncompressed sequencing data, in just 3.5 days [20]. That much data is sufficient to reconstruct up to 12 human-sized genomes, where a single sample of this type equates to a bit over 300 GB of uncompressed data. Moreover, in the context of a study or a sequencing center, this hefty per-sample data size is typically multiplied by a significant number of samples. Consider that for sequencing-based studies that require high analysis sensitivity or to get population-wide statistics, thousands of samples need to be sequenced and processed; e.g., a population-wide study by Orrù et al. [5] required the sequencing of 2870 whole genomes resulting in almost one petabyte of sequencing data to be analyzed and stored.

### A. BCL conversion and sorting reads

The first step in making sense of the raw data is to reconstruct the original DNA fragments from the “slices” produced by the sequencer. The operation required to reconstruct the reads is logically equivalent to a matrix transposition, where a read is obtained by concatenating the elements located at the same positions across several files, as illustrated in Fig. 1.

**Fig. 1:** Recomposing nucleotide sequences from the raw data in Illumina BCL files. The operation requires reading data simultaneously from a number of files equal to the number of cycles in the sequencing run – each file contains one base from each read. Once all the bases of a read are acquired, the read can be composed and emitted and processing can advance to the next one.

Along with the read reconstruction, one must typically also perform other operations which help the subsequent analyses: filtering out some of the reads (based on filter files which are also part of the run output) and tagging each read with some metadata (e.g., location of read within the tile).

Moreover, for reasons of efficiency and flexibility, the DNA fragments from individual samples are often tagged with a short identifying nucleotide sequence and then mixed and sequenced with other samples in a single batch. This procedure is known as *multiplexed sequencing*. In this way, the sequencing capacity of the run is divided among more biological samples than would be possible if it was necessary to keep them physically separate. Being that the genetic material of multiple samples is in the same biological solution, the sequencer will acquire their fragments of DNA indiscriminately and they will be output in the same dataset. Thus, after BCL conversion, the NGS data will typically undergo a *demultiplexing* stage, where fragments from various samples are sorted into separate datasets using their identifying tag.

### B. Read Alignment

The product of the high-throughput shotgun sequencing and processing procedure described thus far is a set of billions unordered short DNA fragments. These need to be analyzed computationally to understand the structure of the original whole DNA sequence from which they came. The specific analyses to be performed will vary depending on the application. The most common sequencing scenario – and the one that concerns us in this work – is the one where there is a reference genome available for the sample being sequenced and it is used as a guide to reconstruct the sample’s genome. This type of sequencing experiment is known as *resequencing*. When resequencing, after the reconstruction of the sequenced fragments, the fragments are *aligned* or *mapped* to the reference genome – i.e., an approximate string matching algorithm is applied to find the position in the reference sequence from which the reads were most likely acquired. The mapping algorithms are designed to be robust to sequencing errors and to the small genetic variations that characterize the sample. Moreover, when sequenced the genome is typically oversampled many times; for instance, for whole-genome sequencing 30 times oversampling – also known as *30X coverage* is common. The oversampling results in reads that overlap, which will be essential in later phases of analysis to detect heterozygosity and to distinguish genomic variants specific to the sample from mere sequencing errors.

### C. Standard practice

In a typical scenario, after the sequencer finishes its work a multi-step processing pipeline will be executed, starting with the BCL conversion and read alignment steps described in the previous subsections. In the state-of-the-art, sequencing operations are equipped with a conventional HPC cluster with a shared parallel storage system. The nodes are typically accessed via a batch queuing system, and often the computing resources are dedicated to the needs of the sequencing and bioinformatics work [12].

Within this context, the standard solution is to perform read reconstruction and demultiplexing using Illumina’s own proprietary, open-source tool: bcl2fastq2. It is written in C++ and powered by the Boost library [23]. This tool implements shared-memory parallelism – i.e., it only exploits parallelism within a single computing node. To the best of the authors’ knowledge, there are no alternatives for this tool in a conventional computing setting.

On the other hand, there is variety of alignment programs available and in widespread use [24], [25]. Among these, BWA-MEM [26] is quite popular and has been found to produce some of the best alignments [27], [28]. Like the bcl2fastq2 – and the other conventional read alignment programs – BWA-MEM also implements shared-memory parallelism.

The output of BWA-MEM is written as Sequence Alignment Map (SAM) files [29]. SAM is a textual format, and hence rather lengthy. Typically, SAM files are then converted to more space-efficient formats, like BAM [29] or CRAM [30], using SAMtools [31]. The CRAM format uses reference-based compression to perform efficient lossless data size reduction and it supports, in a flexible and extendable way, a variety of lossy compression algorithms. In this paper we adopt CRAM as output format and the reference workflow we are considering is one that takes raw NGS data as its input and produces CRAM compressed datasets ready to be further processed by, e.g., variant calling steps.

Though shared-memory parallelism is certainly beneficial to accelerating analysis, it is insufficient. Because of the huge amount of data produced by current sequencers the processing of a run on a single node easily takes several hours of computation, even on state-of-the-art machines, with tens or even hundreds of cores. One way to overcome this problem is to distribute the data among different nodes and then running distinct instances of the programs on each node, each one working on its subset of the data. However, this ad hoc solution is not trivial to implement properly, as one would inevitably end up trying to reimplement one of the already existing distributed computing frameworks.

## III. BACKGROUND

### A. Apache Flink

The processing of big data sets often involves the application of different algorithmic steps, such as filtering, selecting, pre-processing and so on. A traditional way of allowing communications between the different steps while maintaining their independence is by handling the communication via intermediate files: the output files produced by a step become the input for the next one. This approach typically induces two problems:

1. A step can begin only when the previous one has completely terminated;
2. If some new data are to be included in the computation process, updating the final results might need the whole computation pipeline to be re-executed from scratch.

The stream computing paradigm tries and overcome these problems by establishing a continuous flow of data from the source, to which modules can be attached and detached easily, to build computational workflows that can be updated painlessly.

In recent years there has been a flourishing of stream-oriented big data frameworks, such as, e.g., Spark Streaming [32], Storm [33], Samza [34], Apex [35] and Flink [36] (all of which are Apache projects). We have adopted Apache Flink because of its native streaming capabilities, low latency, exactly-once semantics, powerful computing primitives and clean API.

### B. Apache Kafka

Concurrently with the development of stream computing frameworks, there has been an increasing demand for communication means between the streaming modules that could guarantee low latency, fault-tolerance, scalability and flexibility. Some important projects which try and meet these demands are Amazon Simple Queue Service (SQS) [37], Amazon Kinesis [38] and Apache Kafka [39].

We have chosen to adopt Kafka, which is a message broker based on a publish-subscribe model. In addition to offering the features outlined above, is also well supported by Apache Flink, via a dedicated connector. Furthermore, starting from version 0.11, it also offers exactly-once semantics, thus guaranteeing that, even in case of failures, each record is eventually delivered to its consumers and that no duplicates are created in the process.

## IV. IMPLEMENTATION

Our objective is building scalable, robust and easily composable tools, that enable the processing of continuous streams of data, while still being able to be communicate smoothly with the already existing distributed analysis tools (first of all, the Genome Analysis Toolkit (GATK) [40]). Our pipeline is free software and will soon be released to the public.

### A. Data preprocessing

The first module in our pipeline takes care of preprocessing the raw Illumina data, which are available in the proprietary BCL format. To construct the preprocessor we extended our BCL to FASTQ converter [15], by enabling its output to be sent to a Kafka broker, using the built-in Flink-Kafka connector.

Schematically, the preprocessor works as follows:

1. Bases and quality scores are decoded from the cyclebased input files;
2. A data transposition is performed to obtain the reads;
3. The reads are then sorted by their label (i.e., *demultiplexed*);
4. Tiles are processed in parallel, with the degree of parallelism specified as a runtime parameter;
5. For each tile, a list of the new Kafka data topics which will be created is first sent to a special control topic, each new data topic corresponding to a logical output (e.g., LABEL-NAME_L002_2208, where LABEL-NAME is the sample id, while the other two numbers indicate, respectively, the specific lane and tile read);
6. The reads are written into the data topics;
7. Finally, an EOS marker is appended to each data topic.

The preprocessor starts sending the reads to Kafka as soon as they are produced, i.e., to initiate the communication it doesn’t wait for the whole tile to be finished. Similarly, the data topics are created incrementally while the input files are processed. In this way modules which are interested in consuming the topics can start processing as soon as the data are produced, without introducing additional latencies, either between or within topics, as opposed to what happens in the standard pipeline.

### B. Alignment

The alignment module, implemented from scratch in Flink for this work, exploits our Read Aligner API (RAPI [15]), which in turn relies on a modified version of the standard BWA-MEM aligner [26]. The module consumes the reads via TCP from the Kakfa broker (which could thus be located far from the computation nodes used in this step) and it produces as output the aligned reads. Since we are testing a data compression pipeline consisting of two modules, we write the output as CRAM files in an HDFS filesystem, but the program also allows to continue the stream, by sending the output to a (possibly different) Kafka broker.

This module works as follows:

1. The control topic is polled every 3 seconds, to receive notice of newly opened data topics;
2. The data topics are grouped into Flink jobs and opened in parallel, with the degree of parallelism specified as a runtime parameter;
3. For each data topic, the reads are passed to the RAPI/BWA-MEM library in chunks of a few thousands and aligned;
4. The aligned reads are then sent to the output sink, i.e., written as CRAM files using the Hadoop-BAM libraries [41].

## V. EVALUATION

We evaluated the performance and scalability of our Flink-Kafka pipeline by testing it on a real humane genome dataset, running on the Amazon Elastic Compute Cloud (EC2 – https://aws.amazon.com/ec2) and comparing it against the standard pipeline. The software packages we used are detailed in Table II.

**TABLE I:** Maximum sequencing capacities for a number of Illumina high-throughput sequencers [20]–[22]. The size of the output in GB is intended for uncompressed reads with base qualities and id strings, considering a total of 2.5 bytes/base.

| Sequencer | Read length | Time (days) | Gigabases | Gigabytes (GB) | Rate (GB/day) |
| --- | --- | --- | --- | --- | --- |
| HiSeq 2000 | 2 x 100 | 8 | 400 | 1000 | 125 |
| HiSeq 2500 | 2 x 125 | 6 | 1000 | 2500 | 417 |
| HiSeq 4000 | 2 x 150 | 3.5 | 1500 | 3750 | 1071 |

**TABLE II:** Versions of the software packages used in this work.

| Package | Version | Source |
| --- | --- | --- |
| bcl2fastq2 | v2.17.1.14 | http://support.illumina.com/ |
| BWA | 0.7.8-r455 | https://github.com/lh3/bwa |
| Flink | 1.3.1 | https://flink.apache.org |
| Hadoop | 2.7.3 | http://hadoop.apache.org/ |
| Java | Oracle JDK 8u131 | http://www.oracle.com/ |
| Kafka | 0.10.2.1 | https://kafka.apache.org |
| Rapi | Commit ID: 605a5a8 | https://github.com/crs4/rapi |
| SAMtools | v1.5 | https://github.com/samtools/samtools |

### A. Hardware

We ran our experiments using up to 12 instances of type i3.8xlarge, whose characteristics are summarized in Table III.

**TABLE III:**
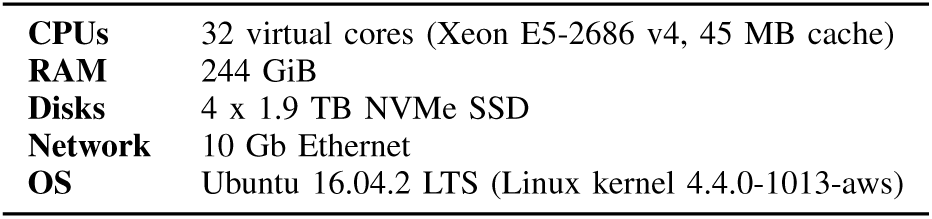
Configuration of the Amazon EC2 i3.8xlarge nodes used to evaluate performance.

### B. Input dataset

The input dataset was produced by an Illumina HiSeq 3000 at the CRS4 Sequencing and Genotyping Platform (at the same research organization as the authors of this manuscript). The run used a single multiplexing tag per fragment and contained 12 human genomic samples, amounting to 47.8 GB of raw data scattered among 47,050 gzip-compressed BCL files plus 224 filter files (which contain a QC pass/fail for each read). The size of each uncompressed BCL and filter file is 4.1 MB. Additional information is reported in Table IV.

**TABLE IV:** Characteristics of the input dataset used for evaluation.

|  |  |
| --- | --- |
| **Sequencer** | Illumina HiSeq 3000 |
| **Size of data files** | 47.8 GB (gzip-compressed) |
| **Number of files** | 47,040 |
| **Number of samples** | 12 |
| **Number of reads** | $3.38 \cdot 10^8$ |
| **Number of DNA bases** | $3.38 \cdot 10^{10}$ |

### C. Experimental Workflow

A straightforward way of running of our pipeline would be

1. Running the pre-processor on some slim nodes, close to the sequencer;
2. Sending the data to a Kafka broker inside a data and compute facility;
3. Running the aligner on a cluster of powerful computing nodes at the facility.

However, to present a fairer comparison with the baseline and to have the same setup for all the configurations tested, we chose to have both preprocessor and aligner running on the same nodes, at the same time. In more detail, to evaluate the speed and scalability of our distributed-memory workflow, we prepared the following setup, on nodes from Node-1 to Node*n*, with 1 *≤ n ≤* 12:

1. A Kafka broker running on Node-1, using all its four SSD disks as buffer space;
2. HDFS distributed among the *n* computing nodes, with each datanode using all its four SSD disks to store the data;
3. The preprocessor running on Flink-on-YARN, on the same *n* nodes;
4. The aligner running within Flink in stand-alone mode (i.e., outside of YARN), on all *n* nodes;

*1)Preprocessing:* To fully test the streaming paradigm we chose to have preprocessing and alignment working concurrently and thus we tuned the parallelism of the preprocessor in order to maintain, for about half of the total runtime, a feed of new data for the aligner (see Table V). To avoid interferences at the scheduler level between preprocessing and alignment, we chose to run the preprocessor inside a dedicated YARN instance, allocating 5 GB of RAM to the Flink Job-Manager and 30 GB to each Task-Manager. The data initially reside in the HDFS from which they are read, processed and sent via TCP to the Kafka broker, running on Node-1.

**TABLE V:** Configuration parameters of the preprocessing module.

|  |  |
| --- | --- |
| **Dedicated RAM per node** | 30 GB |
| **Flink jobs per node** | 4 |

*2)Alignment:* We ran the alignment module in Flink in stand-alone mode, i.e., outside of YARN, and independently from the preprocessing phase. To limit the CPU time wasted in setting up new jobs, we gathered 4 data topics into the same Flink job and set an available parallelism larger than the number of cores, so that when the system is idle loading a new job, it can still work on the other ones (the program parameters are shown in Table VI). We dedicated 10 GB of RAM to the Job-Manager and 180 GB to each Task-Manager.

**TABLE VI:** Configuration parameters of the aligning module.

|  |  |
| --- | --- |
| **Dedicated RAM per node** | 180 GB |
| **Flink jobs per node** | 8 |
| **Data topics per Flink job** | 4 |
| **RAPI parallelism per data topic** | 4 |
| **RAPI window size** | 3360 |

### D. Baseline workflow

As a baseline for comparison we implemented a full standard workflow which handles conversion from BCL to FASTQ, alignment and finally conversion from SAM to CRAM, using the tools outlined in Table II. As described in Section II-C the standard practice only exploits shared-memory parallelism, and hence the baseline was only tested on a single node.

The workflow can be summarized as follows:

1. Conversion from BCL to FASTQ, using the proprietary Illumina program bcl2fastq2;
2. Alignment with BWA-MEM, reading the FASTQ files and writing the output to FIFO pipes;
3. Conversion from SAM to CRAM using SAMtools, reading from the FIFO pipes and writing to the local filesystem.

The exact scripts used to measure the runtime of the baseline are available at https://github.com/crs4/2017-flink-pipeline.

## VI. RESULTS & DISCUSSION

### A. Full pipeline

The runtimes obtained by the experiments are shown in Table VII. On a single node our pipeline is 11% slower than the baseline, the slowdown accounting for the overheads caused by the Flink scheduler and the communication layer. However, as the number of nodes increases, our pipeline achieves near optimal scalability, as shown in Figure 2. The relative scalability (i.e., compared to our runtime on a single node) is 10.6 on 12 nodes, whereas the absolute one (i.e., compared to the faster baseline) is 9.5 on 12 nodes. This result is particularly remarkable since the runtime on 12 nodes is below 15 minutes, and the constant costs of running Flink, YARN and Kafka begin to have a noticeable impact on the total runtime.

**Fig. 2:**
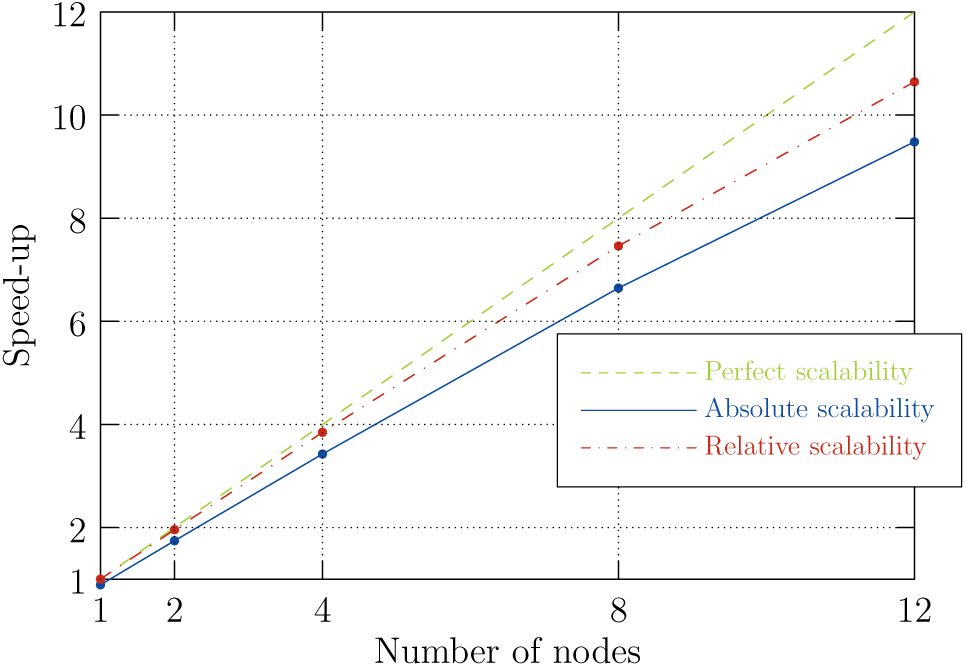
Strong scaling of our Flink/Kafka pipeline, compared with the single-node baseline.

**TABLE VII:** Running times of our Flink/Kafka pipeline and the baseline on a single node.

| Nodes | Time (minutes) |
| --- | --- |
| Baseline | 137 |
| 1 | 152 |
| 2 | 77 |
| 4 | 39.6 |
| 8 | 20.4 |
| 16 | 14.3 |

### B. Kafka

In addition to the scalability experiments, we also tested Kafka to see how it behaves under stress and how many resources it consumes. To this purpose we setup the following 4 nodes configuration:

- Node-1: Running the Kafka broker and Flink Job-Manager;

- Node-[2-4]: Running the preprocessor at full speed, i.e., with 32 concurrent jobs per node.

The above configuration produces a flow of 450 MB/s of data towards the broker for about 3 minutes, creating 2688 topics and filling them with about 340 million messages. The CPU usage to handle this load was about 900%, i.e., 9 cores running at full load. Assuming linearity, this means that when the network bandwidth is saturated (i.e., at 10 Gb/s = 1250 MB/s) the broker will use about 25 cores. As for the RAM usage, Kafka is not memory-greedy and its RAM usage was comfortably below 10 GB.

### C. Limits to scalability

The minimum throughput required to sustain an aligning computing node at full computation power is about 10 MB/s (which in turn gives back an output from the aligner to the broker of about 6.5 MB/s). We would need 125 nodes to saturate the (full-duplex) network bandwidth of the Kafka broker at this data rate, which provides us with an upper bound for the scalability of the cluster setup used in our experiments.

Since the network becomes our bottleneck when the cluster size increases, if more computation nodes are required to work in parallel, one could easily add more Kafka brokers to the configuration, to have the load automatically distributed among them and thus allowing the cluster to grow (at most) additional 125 nodes per each new Kafka broker.

## VII. RELATED WORK

The development of bioinformatics tools based on “Big-Data” technologies started in late 2008 with work on Hadoop [42], [43] and has since continued to increase [44]. Most of the advancement in this area have come in NGS data analysis, where many Hadoop-based tools have been developed [45], [46], [46]–[48] and, more recently, [13], [17]. On the other hand, there is very little on distributed streaming computing applied to the life-sciences, a part from general architecture descriptions such as [49].

## VIII. CONCLUSION

We have presented a fully scalable streaming pipeline to process raw Illumina NGS data up to the stage of aligning reads – the first, to the best of the authors’ knowledge, example of using a distributed publish-subscribe messaging system (Kafka) to decouple between processing steps of a bioinformatics pipeline. Although the practical application considered here is the direct compression of raw NGS data to the CRAM format, by itself a useful task, the program also allows to continue the stream, by sending the output to a (possibly different) Kafka broker and further processing steps that could, for instance, continuously aggregate results across multiple sequencing runs.

Our experiments have shown the pipeline to have very good scalability characteristics, so that a NGS production facility could reasonably expect to reduce their processing per sequencing run to under an hour using a small commodity cluster. Although scalability has been shown up to 12 nodes, there is circumstantial evidence that, in the chosen cluster setup and application, a single Kafka broker will be able to support scaling up to about one hundred nodes.

The code presented it this work will soon be available, as free software at http://github.com/crs4.

## ACKNOWLEDGMENTS

We thank Gianmauro Cuccuru for the NGS dataset used in the experiments, Massimo Gaggero for his support in setting up the AWS EC2 environment and Brendan Lawlor for interesting discussions on Kafka and its potential uses.

